# Genetic similarity enhances the strength of the relationship between gut bacteria and host DNA methylation

**DOI:** 10.1101/2021.07.10.451923

**Authors:** Jia Zhou, Kiflu Tesfamicael, Shao Jia Zhou, Lee A. Rollins, Carlos M. Rodriguez Lopez

## Abstract

Factors such as host age, sex, diet, health status and genotype constitute the environmental envelope shaping microbial communities in the host’s gut. It has also been proposed that gut microbiota may be influenced by host epigenetics. Although the relationship between the host’s genotype/epigenotype and its associated microbiota has been the focus of a number of recent studies, the relative importance of these interactions and their biological relevance are still poorly understood. We used methylation-sensitive genotyping by sequencing to genotype and epigenotype invasive cane toads (*Rhinella marina*) from the species’ Australian range-core (three sites) and the invasion-front (three sites), and 16S rRNA gene sequencing to characterize their gut bacterial communities. We tested the effect of host genotype and epigenotype (i.e., methylome) on gut bacterial communities. Our results indicate that the genotypes, epigenotypes and gut communities of the range-core and invasion-front are significantly different from one another. We found a positive association between host pairwise genetic and epigenetic distances. More importantly, a positive relationship was found between the host’s epigenetic and gut bacterial pairwise distances. Interestingly, this association was stronger in individuals with low genetic differentiation. Our findings suggest that in range-expanding populations, where individuals are often genetically similar, the interaction between gut bacterial communities and host methylome may provide a mechanism through which invaders increase the plasticity of their response to novel environments, potentially increasing their invasion success.

## Introduction

The gut microbiota can play a key role in host adaptation to the environment by affecting its phenotype (Alberdi et al. 2016). Concurrently, host factors such as age, sex and health status contribute to the environmental envelope shaping gut microbial communities (Pereira et al. 2020; Tong et al. 2020). In addition to these factors, host genetic diversity also may be an important determinant of host-microbial relationships. For example, host heterozygosity (within-individual genetic variation) has been positively associated with individual fitness and adaptive potential (Mainguy et al. 2009; Velando et al. 2015). Such heterozygosity-fitness correlations have been widely studied, including in the context of disease/parasite resistance, and host body mass, reproductive performance and survival (Eastwood et al. 2017; Coltman et al. 1999; Penn et al. 2002; Luikart et al. 2008; Mainguy et al. 2009; Velando et al. 2015; Brambilla et al. 2018). In gut microbial studies, alpha diversity (microbial diversity within individual hosts) also has been associated with increased host fitness (e.g. resistance to parasites and disease; Kreisinger et al. 2015; Estaki et al. 2016; Suzuki 2017). These results suggest a positive correlation between host genetic diversity and microbial alpha diversity. However, a negative relationship between these metrics has been found in at least one species (fur seals; Grosser et al., 2019), indicating that further analysis of these relationships in a broader range of taxa would allow a better understanding of how the host’s genetic diversity affects gut microbial community diversity.

In addition to the degree of host genetic diversity within an individual, the patterns of genetic variation across the genome warrant investigation with respect to interactions with the host’s microbial community. Even though gut microbiota is largely acquired from the environment (Alberdi et al. 2016), this community also can be shaped by host genotype (Goodrich et al. 2014; Blekhman et al. 2015; Goodrich et al. 2016). Particular host genotypes have accounted for substantial differences in microbial community composition and diversity; for example, in humans, microbiota variation was driven by immunity-related host genotype (Blekhman et al. 2015). This suggests that there could be a heritable component to gut microbial composition. Microbiome composition of desert bighorn sheep (*Ovis canadensis nelsoni*) diverged in accordance with both landscape-scale environmental and host population characteristics (Couch et al. 2020). Stickleback gut microbiota variation across populations was associated with host genotype more than with environmental factors (Smith et al. 2015). Conversely, host genetic effects were much weaker than the environment in shaping the human gut microbiota (Rothschild et al. 2018). Collectedly, these results indicate that the relative strength of host genetic versus environmental influence on gut microbiota may be species-specific.

Gut microbiota may also interact with host epigenotype, providing a mechanism through which gut microbial communities can affect host health and adaptation (Stilling et al. 2014; Krautkramer et al. 2016). For example, clear associations between bacterial composition and host DNA methylation profiles have been identified in relation to body weight and metabolism regulation (Kumar et al. 2014; Cuevas-Sierra et al. 2019). Additionally, gut microbiota guides and/or facilitates epigenetic development of intestinal stem cells during the postnatal period and may influence lifelong gut health (Yu et al. 2015). At the same time, host epigenetic status may affect gut microbiota: DNA methylation in intestinal tissue is known to contribute to the regulation of genes involved in cell proliferation, anti-bacteria metabolite production, anti-inflammation and to play a critical role in re-establishing gut homeostasis in mice (Ansari et al. 2020; Wu et al. 2020). These results suggest that it is important to examine the interaction between gut bacteria and host DNA methylation, and that this relationship can be bi-directional.

An individual’s genotype can strongly influence their epigenotype (Bell et al. 2011; Dubin et al. 2015). Additionally, genetic variation can contribute to the transgenerational heritability of DNA methylation (in humans; McRae et al. 2014). Genetic effects can be stronger than the effects of exogenously manipulated DNA methylation (in cane toads; Sarma et al. 2020). In addition to the relationship between an individual’s genotype and epigenotype, at the population level, an increased variability in DNA methylation may occur in populations with low genetic variation, as compared to populations with higher genetic variation (Liebl et al. 2013). In particular, this has been discussed in the context of expanding range-edge populations of invasive species, and it has been hypothesised that this may facilitate adaptation to new environments by creating phenotypic diversity (Ardura et al. 2017; Sheldon et al. 2018; Hawes et al. 2018; Carja et al. 2017).

Although these relationships between gut microbiota, host genotype and host epigenotype (e.g. DNA methylome) have been examined in humans and domesticated animals (Goodrich et al. 2016; Cuevas-Sierra et al. 2019; David et al. 2019; Ansari et al. 2020; Ryan et al. 2020; Xu et al. 2020), little of this research has been conducted in non-model species. Further, the relevance of these relationships to invasion success is virtually unexplored. Here, we used the iconic invasive cane toad to conduct the first characterisation of these relationships in an amphibian and to determine whether these relationships change when comparing samples collected across an expanding invasive range. Although gut bacterial communities and toad genetics have been previously characterised across Australia, their relationship to each other has not been studied. We have previously found significant differences between the gut bacterial communities of range-core and invasion-front cane toads (Zhou et al. 2020). Host genetics also differ across the range: population structure has been identified across Australia and genetic diversity is reduced at the range edge (Selechnik, Richardson, Shine, DeVore, et al. 2019). Moreover, substantial shifts in gene expression in spleen and muscle tissue were identified between invasion-front and range-core toads (Rollins et al. 2015; Selechnik, Richardson, Shine, Brown, et al. 2019). To date, there have been no investigations of DNA methylation in wild-collected toads in Australia.

We investigated whether the variation observed in cane toad gut bacteria across Australia is mediated by host genetics, and whether gut bacterial communities are correlated with host DNA methylation. Specifically, we tested the following hypotheses: (1) if host heterozygosity and diversity in gut bacteria are both positively associated with individual fitness and long-term population persistence, then more heterozygous hosts will have higher gut bacterial diversity; (2) if epigenetic diversity acts as a compensatory factor in populations with low genetic diversity, then genetic diversity will be negatively correlated with DNA methylation diversity; (3) if the host genotype is an important contributor to the environmental envelope in which gut microbes develop, then genetically similar toads will share similar gut bacteria; and (4) if gut bacteria can cause heritable phenotypic changes through epigenetic modification and DNA methylation in intestinal tissue is known to play a critical role in re-establishing gut homeostasis, then cane toads that possess similar gut bacteria will also have similar DNA methylation profiles.

## Results

### Characterisation of host genetic, host DNA methylation and gut bacterial diversity and differentiation

A total of 55 individuals were used for all datasets analysis, after removing those whose msGBS reads from blood samples presenting low coverage. We first investigated within-individual genetic and gut bacterial community by calculating individual heterozygosity (HL) and gut bacterial alpha diversity (Shannon’s diversity Index). The observed mean HL was 0.49 (SD: 0.03) and the mean of Shannon’s index was 6.00 (SD = 0.92).

Principal coordinate analysis (PCoA) plots showed host SNP, DNA methylation and gut bacterial profiles clustered according to their provenance, first by sampling site and then by region (invasion-front or range-core) (Figure 1 A, B, and C, respectively). PCoA plots suggested similar levels of diversity for all three profile types. However, comparison of Bray Curtis values from SNP data indicated that range-core toads had significantly higher levels of genetic diversity than those from the invasion-front (invasion-front: mean = 0.126, SD = 0.009; range-core: mean = 0.140, SD = 0.010; t test: t = 19.569, df = 710.3, p-value < 0.01); while range-core and invasion-front toads had similar levels of diversity of DNA methylation (invasion-front: mean = 0.121, SD = 0.022; range-core: mean = 0.122, SD = 0.019; t test: t = 0.232, df = 723.42, p-value = 0.82) and gut bacterial community diversity (invasion-front: mean = 0.640, SD = 0.132; range-core: mean = 0.656, SD = 0.114; t test: t = 1.708, df = 723.19, p-value = 0.09). Genetic, epigenetic and gut bacterial community pairwise comparisons show significant differences in beta diversity (diversity between samples) between most sampling sites (Table 1). Finally, we found significant differences between invasion-front and range-core sampling sites in host genotypes (adonis2: R^2^ = 0.089, F = 5.162, p-value < 0.001; betadisper: F-value = 23.337, p-value < 0.001), host epigenotypes (adonis2: R^2^ = 0.104, F = 6.120, p-value < 0.001; betadisper: F-value = 0.005, p-value = 0.942) and gut bacterial community (adonis2: R^2^ = 0.099, F = 5.830, p-value < 0.001; betadisper: F-value = 0.226, p-value = 0.636).

**Table 1.**
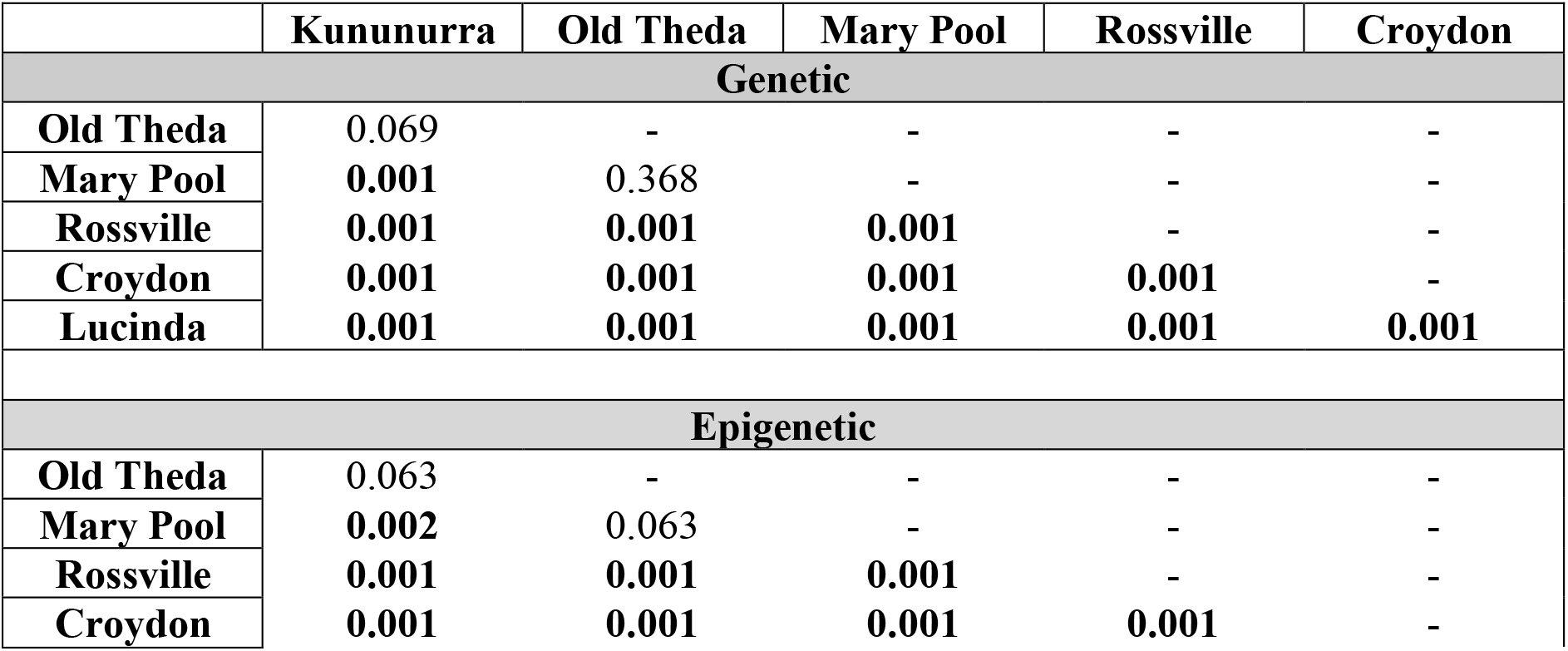
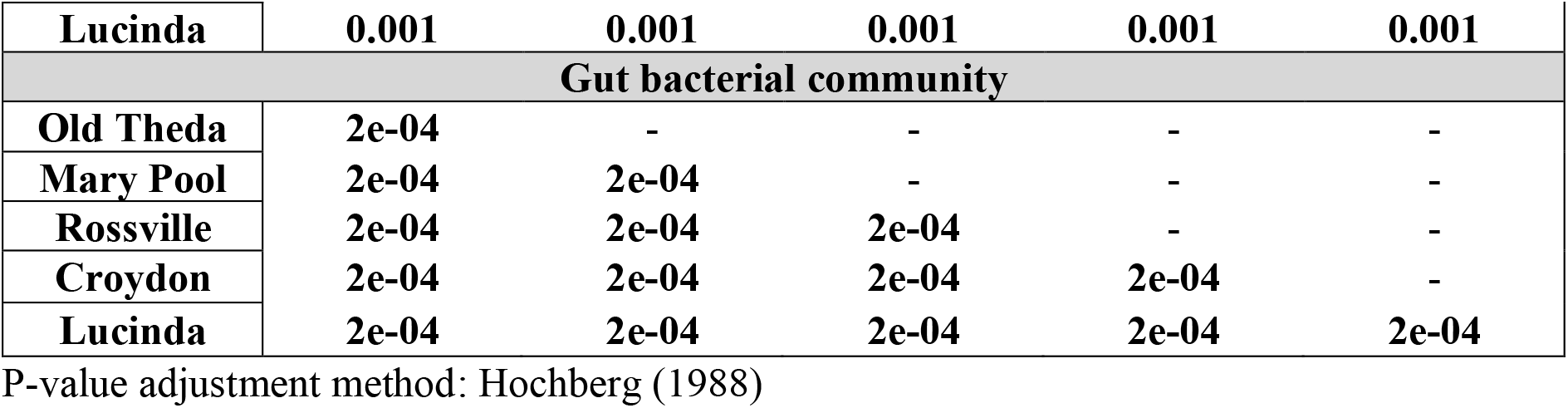
Analysis of genetic, epigenetic, and gut bacteria differences between cane toad sampling sites in the Australian invasion-front and range-core. Pairwise comparisons were calculated using permutation MANOVAs on Bray Curtis genetic, epigenetic, and gut bacterial community distance matrices from 10 female cane toads per sampling site in the invasion-front (Kununurra, Old Theda, Mary Pool) and range-core (Rossville, Croydon, Lucinda). Genetic and epigenetic distances were estimated based on SNPs and DNA methylation epialleles identified using msGBS profiles. Bacterial community distances were calculated using the Core50 gut bacteria taxa identified using 16S rRNA gene sequencing. Bold font indicates significantly different comparisons.

**Figure 1.**
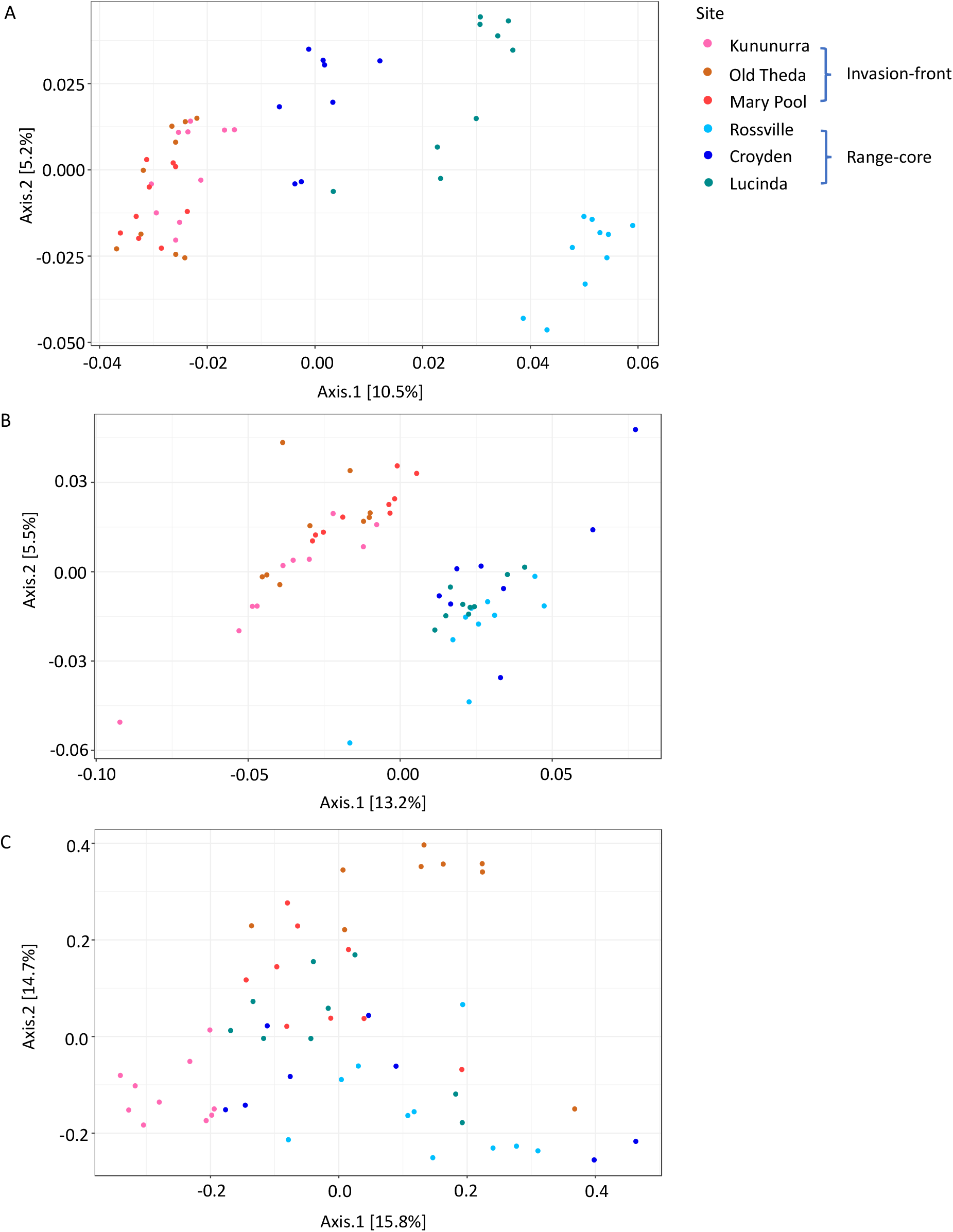
Genetic, epigenetic and gut bacteria differences between wild cane toads in the Australian invasion-front and range-core. Principal coordinate analysis plots were built using Bray Curtis genetic (A), epigenetic (B), and gut bacterial community (C) distance matrices between 6 sampling sites located in the toad’s invasion-front (Western Australian: Kununurra, Old Theda, Mary Pool) and range-core (Queensland: Rossville, Croydon, Lucinda). Genetic and epigenetic distances were estimated based on SNPs and DNA methylation epialleles identified using msGBS profiles. Bacterial community distances were calculated using the Core50 gut bacteria taxa identified using 16S rRNA gene sequencing. All analyses were performed on data collected from 10 female cane toads per sampling site.

### Association between host genetic/epigenetic and gut bacterial diversity and differentiation

Linear mixed model (LMM) analysis accounting for sampling site as a random effect did not identify a correlation between within-individual heterozygosity index (HL) and within-individual gut bacterial alpha diversity (Shannon’s diversity Index) (df = 52.716, t-value = 1.444, p-value = 0.16).

Mantel tests comparing between-individual genetic and gut bacterial community differences showed that genetically similar toads did not share similar gut bacteria (Spearman correlation: Mantel r = 0.0788, p-value = 0.116). On the contrary, toads with similar DNA methylation profiles shared similar gut microbial composition (Spearman correlation: Mantel r = 0.1553, p-value = 0.03). Host genotype and host DNA methylation dissimilarity matrices were positively associated (Spearman correlation: Mantel r = 0.654, p-value = 0.001).

LMM analysis with gut microbial distance as the response variable indicated that gut bacterial differentiation between individuals is affected by: 1) host DNA methylation differentiation (df = 1461.645, t-value = 2.505, p-value = 0.01), and 2) the interaction of host genetic distance with host methylation distance (df = 1441.646, t-value = -2.155, p-value = 0.03; Figure 2). The observed relationship between gut bacterial distance and host DNA methylation distance was stronger in cane toad pairs that were more genetically similar (Figure 2). When host DNA methylation distance was used as the response variable, LMM analysis indicated that DNA methylation distance was not affected by gut bacterial distance (df = 1390, t-value = 0.598, p-value = 0.55; Figure 3). DNA methylation distance was significantly associated with genetic distance (df = 1390, t-value = 5.734, p-value < 0.001; Figure 3). There was no interaction between these relationships and population (invasion-front *vs*. range-core).

**Figure 2.**
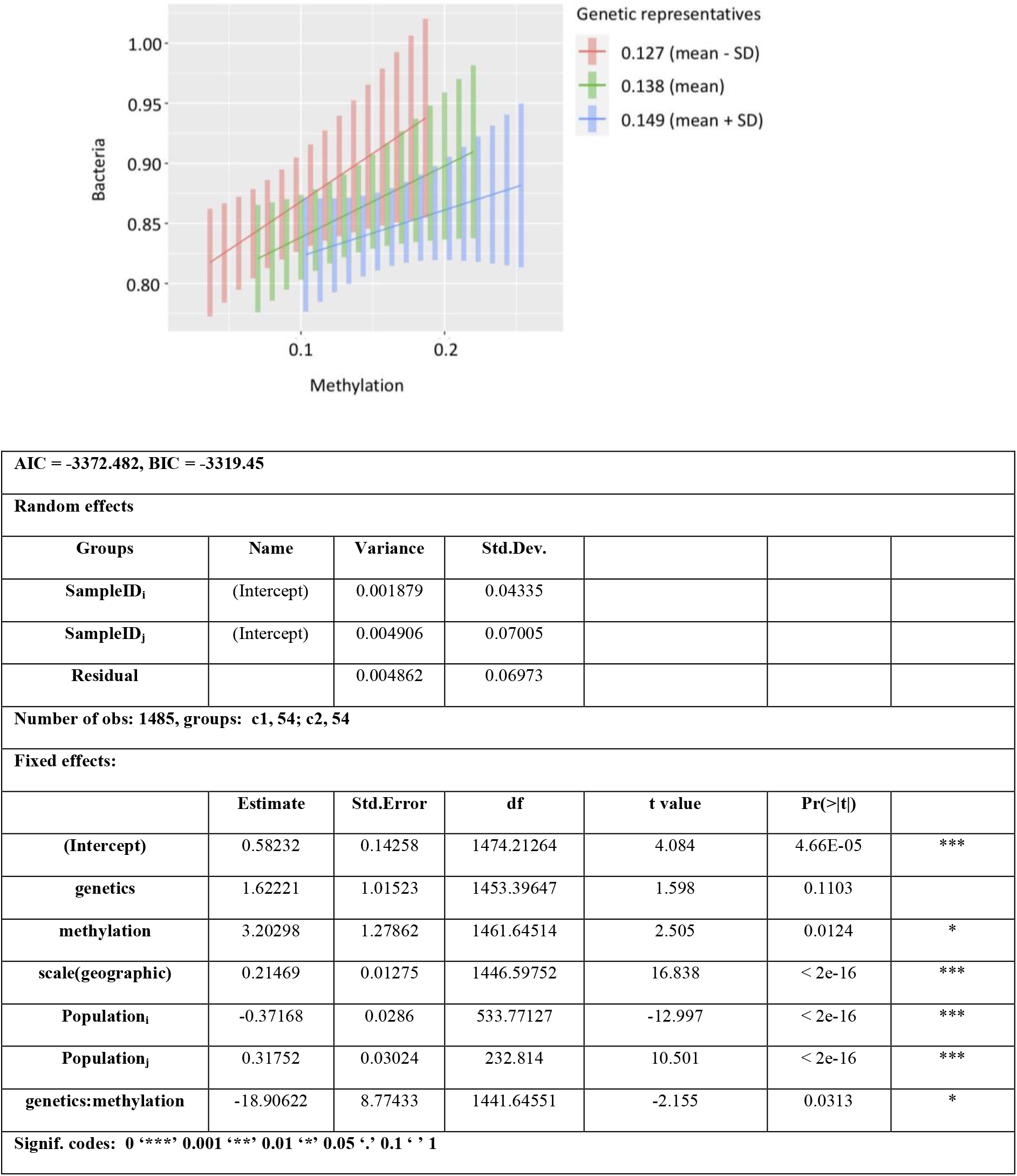
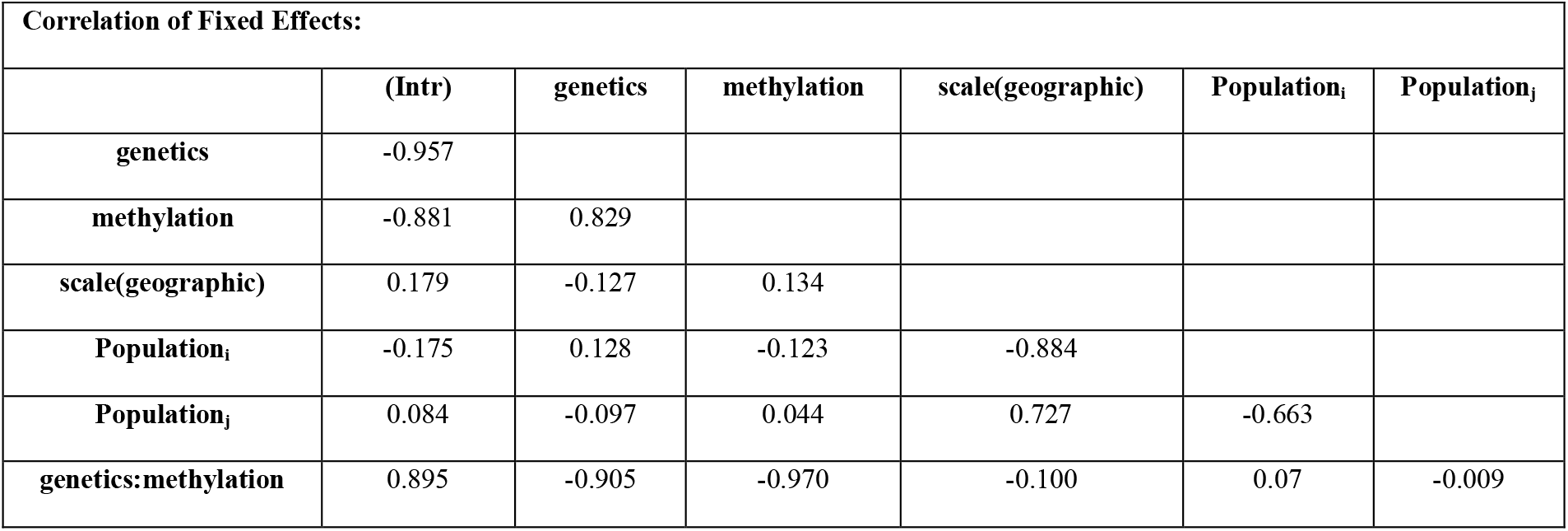
Linear mixed model analysis of the association between gut bacterial community and epigenetic differences among individuals. LMM was based on the Bray Curtis pairwise distances: *Bacteria_dist*_*ij*_ *∼ Genetics_dist*_*ij*_ *\* Methylation_Dist*_*ij*_ *+ scale(Geographic Distance*_*ij*_*) + Population*_*i*_ *+ Population*_*j*_ *+ (1*|*SampleID*_*i*_*) + (1*|*SampleID*_*j*_). Bacterial community distance was considered as the response variable, and genetic, epigenetic, and geographic distance (rescaled), and population (invasion-front or range-core) were considered as fixed factors. Sample IDs as random factor. The figure shows the correlation between epigenetic and bacterial community distances (X axis: fixed factor, Y axis: response factor). Three genetic representative values represent an infinite set of values with which to fix the continuous genetic distances (UCLA: Statistical Consulting Group): the mean level of genetic distance, one standard deviation above the mean level of genetic distance, and one standard deviation below the mean level of genetic distance. LMM model statistics output is shown in the table.

**Figure 3.**
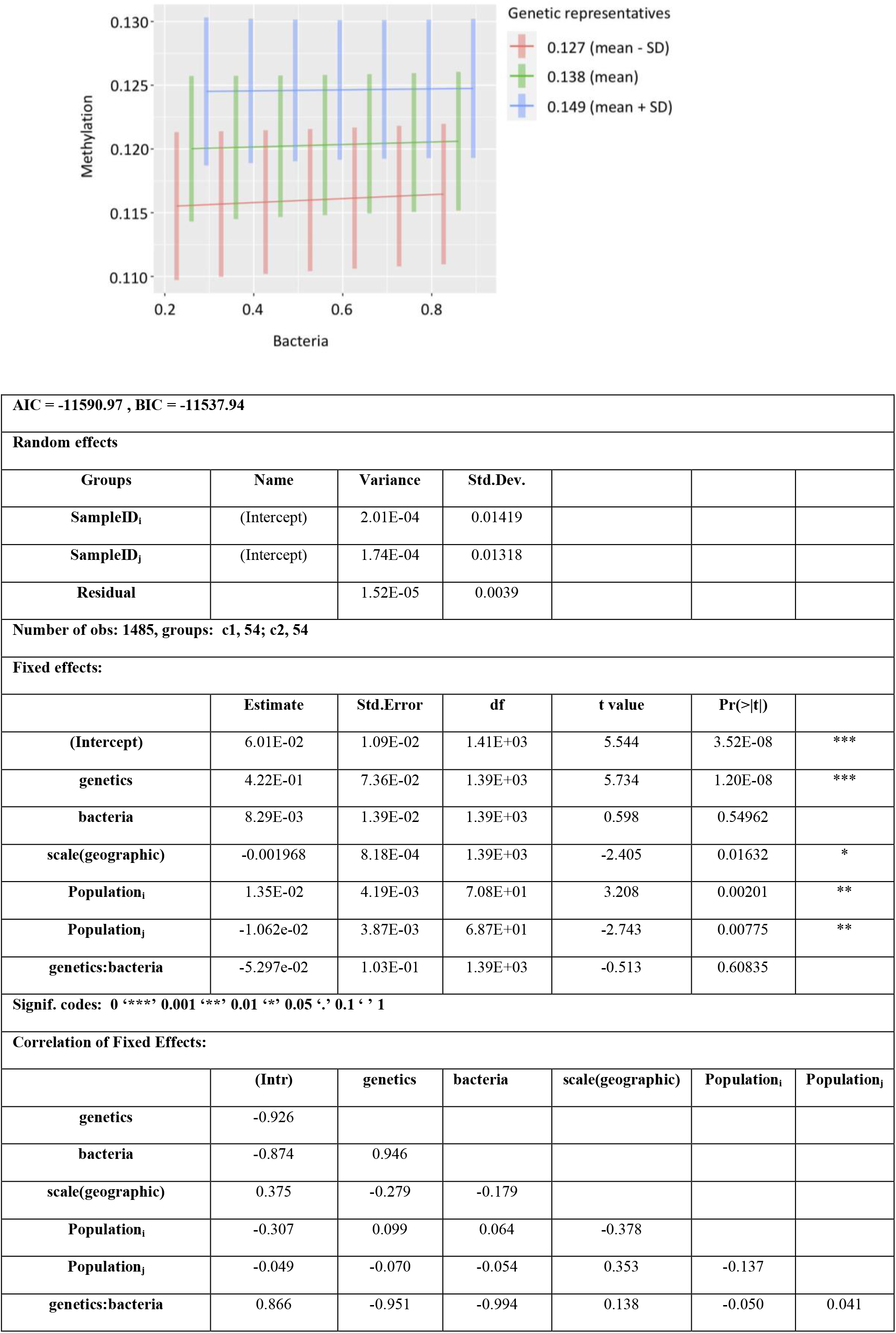
Linear mixed model analysis of the association between epigenetic and gut bacterial community differences among individuals. LMM was based on the Bray Curtis pairwise distances: *Methylation_Dist*_*ij*_ *∼ Genetics_dist*_*ij*_ *\* Bacteria_dist*_*ij*_ *+ scale(Geographic Distance*_*ij*_*) + Population*_*i*_ *+ Population*_*j*_ *+ (1*|*SampleID*_*i*_*) + (1*|*SampleID*_*j*_). Epigenetic distance was considered as the response variable, and genetic, bacterial community, and geographic distance (rescaled), and population (invasion-front or range-core) were considered as fixed factors. Sample IDs as random factor. The figure shows the correlation between epigenetic and bacterial community distances (X axis: fixed factor, Y axis: response factor). Three genetic representative values represent an infinite set of values with which to fix the continuous genetic distances (UCLA: Statistical Consulting Group): the mean level of genetic distance, one standard deviation above the mean level of genetic distance, and one standard deviation below the mean level of genetic distance. LMM model statistics output is shown in the table.

## Discussion

In this study, we used next-generation sequencing to characterize and explore the relationship between genotype, epigenotype (DNA methylation), and gut bacterial communities in wild cane toads across their Australian invasive range. We found no relationship between within-individual genetic diversity and the diversity of its gut bacteria. Additionally, we found that while genetic differentiation was positively related to DNA methylation differences between individuals, there did not appear to be a relationship between the diversity of these two measures. We also found that pairwise differentiation between cane toad gut bacteria was associated with pairwise differentiation between host DNA methylation, and this association was stronger in pairs that were more genetically similar.

There is a growing interest in understanding how host and environmental factors contribute to gut microbiota variation, and how these may impact host phenotype (Zhang et al. 2010; Ussar et al. 2015; Kreznar et al. 2017). However, few studies have investigated the relationship between host heterozygosity and gut bacteria. Because both host heterozygosity (Mainguy et al. 2009; Velando et al. 2015) and gut bacterial diversity (Kreisinger et al. 2015; Estaki et al. 2016) have been reported to be positively related to individual fitness, we predicted that hosts with higher levels of heterozygosity would have more diverse gut bacteria. This relationship has been investigated previously in fur seals (Grosser et al., 2018) and that study found that an individual’s heterozygosity (calculated with microsatellite data) was negatively associated with its microbial diversity. Grosser et al., (2018) proposed that higher quality individuals (who have greater heterozygosity) should be more effective at suppressing nonbeneficial microbes, thus having less diverse microbiotas (Grosser et al., 2018). A negative relationship between these metrics has also been found in sticklebacks, where individuals with greater heterozygosity at the MHC (Major Histocompatibility class II) had less diverse gut microbiota (Bolnick et al. 2014). In this study, we found no relationship between host individual heterozygosity and bacterial diversity in cane toads. The hypotheses in all these studies depend on a positive relationship between heterozygosity and fitness. However, the validity of studies of heterozygosity-fitness correlations, where small numbers of markers (e.g. microsatellites) have been used, has been challenged because the correlation between estimated heterozygosity and true genome-wide heterozygosity is weak (Dewoody & Dewoody 2005; Forstmeier et al. 2012). The SNP data set used here to calculate heterozygosity was large (>38,000 SNPs) and, thus, may provide a more accurate picture of these relationships.

There could be a heritable influence on gut microbial composition, mediated by host genotype (Goodrich et al. 2014; Blekhman et al. 2015; Goodrich et al. 2016). To investigate this in toads, we tested whether genetic similarity between hosts was related to similarity of their gut bacteria. We found no significant association between these metrics. Results of other studies investigating this question are mixed. In chickens, host genetics played a minor role in shaping the gut microbiota (Wen et al. 2019). However, in wild house mice, gut microbiota dissimilarity was significantly correlated with both host genetic distance and body mass index, but not significantly associated with other factors, including diet, climate and geographic distance (Suzuki et al. 2019). Gut microbiota was also found to be significantly correlated with host genetics in fish and amphibians (Griffiths et al. 2018; Uren Webster et al. 2018). Because gut microbiota can be affected by a wide variety of host and environmental factors, it seems likely the relationship between host genetics and gut bacteria is complex and may vary depending on the strength of other host and environmental factors.

Gut microbiota can cause heritable phenotypic changes through epigenetic modification of host genome (Stilling et al. 2014; Krautkramer et al. 2016; Grieneisen et al. 2020). DNA methylation is an epigenetic mechanism known to play a critical role in re-establishing gut homeostasis in intestinal tissue (Ansari et al. 2020; Wu et al. 2020), suggesting this may be a bi-directional relationship. The observed changes in some phenotypic traits (e.g. behaviour) in cane toads across Australia have been linked to their gut bacteria (Zhou et al. 2020). It is possible that this association could be mediated through shifts in DNA methylation in toads. In this study, we found that both DNA methylation and gut bacteria were significantly different between different sampling localities, and that differentiation of host DNA methylation was positively related to differentiation of gut bacteria between pairs of individuals. Because the direction of the relationship between host DNA methylation and gut bacterial community is unknown, we ran separate LMMs using each of these metrics as the dependent variable. Interestingly, when gut bacterial distance was used as the dependent variable, its relationship to DNA methylation distance was stronger in pairs of individuals that were more genetically similar. This dynamic suggests that in populations with more genetically similar individuals (e.g. invasion-front populations), the relationship between DNA methylation and gut bacteria also may be stronger. On the contrary, when DNA methylation distance was used as the dependent variable, there was no significant relationship with gut bacterial distance. The strong influence of host genotype on DNA methylation may mask any potential influence of gut bacteria on DNA methylation.

It is tempting to speculate that the strengthened relationship between gut bacteria and DNA methylation in cane toads that are genetically similar could facilitate cane toad adaptation to novel environments in Australia. First, gut bacterial variation caused by environmental factors (e.g. food resources) could alter host DNA methylation, leading to beneficial phenotypic changes that increase host fitness (Stilling et al. 2014; Krautkramer et al. 2016; Grieneisen et al. 2020). Second, environmental factors could alter host DNA methylation, which could affect the host’s ability to use local microbes or to maintain a balanced gut bacteria by supressing nonbeneficial microbes (Ansari et al. 2020; Wu et al. 2020). Future studies including, fecal transplantation and methylation manipulation experiments will be needed to illuminate the causal relationship and underlying mechanisms in this system.

During invasions, increased variation in host DNA methylation could be compensatory for low genetic diversity, and facilitate adaptation to novel environments by creating phenotypic diversity (Ardura et al. 2017; Carja et al. 2017; Sheldon et al. 2018; Hawes et al. 2018). This is an intriguing idea, and could explain the multitude of phenotype shifts seen in toads as they have spread across Australia (Rollins et al. 2016), despite their low genetic diversity, especially at the invasion-front (Lillie et al. 2014; Selechnik, Richardson, Shine, DeVore, et al. 2019). In this study, we found that although genetic diversity differed across the Australian range, DNA methylation patterns did not, suggesting that no such relationship exists in this invasion. Similarly, in a study of invasive house sparrows (*Passer domesticus*) in Australia, no compensatory relationship between genetic diversity and DNA methylation diversity was detected either (Sheldon et al., 2018). In human populations, diversity of DNA methylation mirrored genetic diversity (Carja et al. 2017). Together, this evidence suggests that further research is needed to understand whether these factors interact to promote phenotypic variation on invasion fronts and, if so, whether the strength of this relationship depends on the degree of influence that the genome exerts on the epigenome for a given species.

## Conclusion

Our results demonstrate that gut bacteria of invasive cane toads in Australia is positively correlated with individual DNA methylation profile changes, and this is accentuated when genetic differentiation is low. DNA methylation variation is similar across the invasion, whereas genetic diversity decreases on the invasion front, suggesting no relationship between the diversity of these metrics. However, genetic differentiation and DNA methylation differentiation have a strong, positive association suggesting that genetic composition determines DNA methylation in this species. These findings provide insights into the dynamics between host genotype, epigenotype and gut bacteria in this iconic invasive amphibian. Moreover, this study draws our attention to the complexity of these relationships and how they may shift over an expanding invasion.

## Materials and Methodology

### Animal materials

We hand-captured 60 wild adult female cane toads from three sites in the Australian invasion-front and three sites in the range-core (Figure S1) and euthanized them by injecting tricaine methanesulfonate (MS222) buffered with bicarbonate of soda. We collected blood and colon content by heart puncture and colon dissection respectively, and preserved these samples in 95% ethanol. Samples were frozen at -20°C for storage until we conducted DNA extractions. The University of Adelaide Animal Ethics Committee approved the collection and use of animals for in this research (approval number: S-2018-056).

### Blood Genomic DNA extraction and Methylation-sensitive genotype by sequencing

We extracted genomic DNA from blood using a PureGene Tissue Kit (QIAGEN), following the manufacturer’s protocols. We performed msGBS on blood DNA as described by Kitimu et al. (Kitimu et al. 2015). In addition to the 60 genomic DNA samples, we included a water blank to account for environmental contamination introduced during sequencing library preparation. We used two enzymes, *EcoRI* (cutsite: GAATTC) and *HpaII* (cutsite: CCGG), to generate restriction products. Enzymatic restrictions were performed in a 16 μl mix containing: 1.6 μl Cut Smart Buffer, 0.32 μl *EcoRI-*HF (NEB #R0101 (20,000 units/ml)), 0.64 μl *HpaII* (NEB #R0171S (10,000 units/ml)), and 13.4 μl DNA (10ng/μl). The enzyme digestion reaction was conducted at 37 °C for 2 h and then 65 °C for 20 min for enzyme inactivation.

A set of barcoded adapters with an *HpaII* overhang and a common Y adapter with an *EcoRI* overhang (Table S1) were used for the ligation reaction. Working stocks of barcoded (0.02 μM) and common Y adapters (3 μM) were prepared in advance as described by Poland et al., (2012). The 32 μl ligation reaction was carried out by adding 0.08 μl T4 Ligase (200 U, NEB) and 3.2 μl T4 Ligase buffer (10X, NEB), 8.72 μl water and 4 μl of the working adapter stock to the 16 μl restriction products. Ligation mixes were incubated at 22 °C for 2 h and 65 °C for 20 min. We removed unused adapters and restriction/ligation products smaller than 100 bp using AMPure XP magnetic beads (x0.9 bead/reaction volume to volume ratio). The clean-up products were used for PCR amplification. Each 25 μl PCR consisted of 10ul digested/ligated DNA library (<1,000ng), 12.5 µl of Q5 MasterMix (Q5 High-Fidelity 2X Master Mix), 2 µl forward and reverse primers @10 µM (Table S1) and 0.5 µl of water. Reactions were performed at 98 °C for 30 sec, 12 cycles for (98 °C for 30 sec, 62 °C for 20 sec, 72 °C for 30 sec) and 72 °C for 5 min. PCR product concentrations were estimated using NanoDrop One^c^ spectrophotometer. Samples were then equimolarly mixed into a single pool. The resultant pool was then split into four subsamples. Fragments below 100 bp and above 600 bp were removed using a magnetic beads and double size selection (x1 bead/reaction volume to volume ratio followed by x0.55 bead/reaction volume to volume ratio). All four size selected fractions were then pooled and quality checked using a Qubit 4 Fluorometer (Thermo Fisher Scientific, Waltham, MA, USA) and a Fragment Analyzer (Agilent, Santa Clara, CA, USA). Sequencing was performed using HiSeq 4000 150bp PE at Novogene Corporation Inc (Sacramento, CA, USA).

### Bacterial DNA isolation and amplicon sequencing

The bacterial DNA isolation, amplicon library preparation and sequencing were described previously (Zhou et al. 2020). Briefly, we extracted bacterial DNA from colon content using the DNeasy PowerSoil kit (Qiagen), following the manufacturer’s protocols. We performed 16S rRNA gene amplicon sequencing on DNA samples by following guidelines for the Illumina MiSeq System. We included 60 colon DNA extracts, one Zymo isolated DNA standard (D6305, community positive control) and one water blank (PCR negative control). We prepared libraries based on the hypervariable (V3-V4) region of the 16S rRNA gene using primers 341F (5’ – **TCGTCGGCAGCGTCAGATGTGTATAAGAGACAG**CCTACGGGNGGCWGCAG-3’) and 785R (5’-**GTCTCGTGGGCTCGGAGATGTGTATAAGAGACAG**GACTACHVGGGTATCTAA TCC-3’) (Herlemann et al. 2011). Sequencing was conducted on the Illumina MiSeq platform at the Ramaciotti Centre for Genomics (University of New South Wales, Kensington, Sydney).

### Data analysis

#### DNA Methylation profiling

We demultiplexed sequencing data with GBSX v1.3 (Herten et al. 2015) and checked quality using FastQC v0.11.4 (Andrews 2010). We trimmed data using AdapterRemoval v2.2.1 (Schubert et al. 2016) and aligned trimmed data to the cane toad reference genome (*Rhinella marina* PRJEB24695; Edwards et al. 2018) using HISAT2 v2.1.0 (Kim et al. 2015). The water blank had very low QC-passed reads (637 reads) and showed low contamination in library preparation and sequencing process. Non-control samples presented an average of 23,330,961 (+/- 21,763,581) QC passed reads, with a mean GC content of 45.96% (+/- 1.95%) and a mean mapping efficiency of 78.21% (+/- 1.11%). Samples presenting less than 5,000,000 reads were removed from further analysis, resulting in the inclusion of 55 cane toad samples. We used SAMtools (Li et al. 2009) to sort and index bam files and then loaded them into Rstudio (R Core Team 2020). We estimated the methylation status of the sequenced loci using “msgbsR” v1.12.0, an R package developed specifically for msGBS data analysis (Mayne et al. 2018). After removing loci not yielding reads in more than 40% of the toad samples and less than one count per million (CPM) in at least 60% of toads using “edgeR” v3.30.3 (Robinson et al. 2010) in R, a total 165,858 loci were kept for further analysis.

#### SNPs profiling

We used BCFtools v1.9 (Li et al. 2009) for SNP calling. We used VCFtools v0.1.15 (Danecek et al. 2011) filtering to only keep variants that have been successfully genotyped in 60% of individuals, a minimum quality score of 30, and a minor allele count of 3. These were then imported as a vcf output file into Tassel v5.0 (Bradbury et al. 2007). Here, only SNPs with at least 0.05 minor allele frequency (Suzuki et al. 2019) were kept (i.e., 38,140 SNPs). For duplicate positions, only the first SNP record was retained and the final SNPs dataset included 38,129 SNPs.

#### Bacterial community profiling

The filtering and processing of raw 16S rRNA sequences were described previously (Zhou et al. 2020). In summary, sequences were filtered by trimming the first 20 bases and truncating each read to 200 bases (based on sequence base quality score), dereplicating, then merging forward/reverse reads, removing chimeras, and finally generating amplicon sequence variants (ASVs) for downstream analysis through the DADA2 pipeline, implemented in QIIME2 v2020.8 (Callahan et al. 2016; Bolyen et al. 2018). In this ASVs table, reads from colon samples and the positive control ranged between 103,245 and 245,059 counts; the PCR negative control yielded 6,727 reads. We used Greengenes version 13_8 to assign taxonomy to the ASVs (DeSantis et al. 2006).

We imported ASVs into the R package “phyloseq” (McMurdie & Holmes 2013) to remove representatives classified to Archaea (n = 28), chloroplast (n = 17), mitochondria (n = 186), and 151 unassigned (“kingdom”) ASVs. We also removed the ASVs with prevalence of less than four, which makes the logged counts per sample more evenly distributed. The remaining 9,878 taxa were classified to the kingdom with 62.62% assigned to phylum level and 39.65% assigned to family level. We imported the pruned ASVs into QIIME2 and calculated observed ASVs (DeSantis et al. 2006), evenness (Pielou 1966), and Shannon (Shannon 1948) indices for bacterial alpha diversity (a measure of microbial diversity within individual host).

We calculated Core50 gut community (Bletz et al. 2016) by filtering ASVs and keeping only those presented in a minimum of 50% of individual toads from each site. This calculation was performed separately for three sites from invasion-front toads: Kununurra (gut Core50: n = 111 ASVs), Old Theda (gut Core50: n = 118 ASVs), and Mary Pool (gut Core50: n = 129 ASVs); three sites from range-core toads: Rossville (gut Core50: n = 148 ASVs), Croydon (gut Core50: n = 86 ASVs), and Lucinda (gut Core50: n = 117 ASVs). We then compiled filtered ASVs of the six sites to avoid excluding ASVs that may be specific to only one site. In combination, the gut Core50 contained 325 unique ASVs, which We used for analysis of beta diversity.

#### Association analysis of heterozygosity and gut bacterial diversity within individuals

Since host heterozygosity and homozygosity matrixes are highly correlated (Charpentier et al. 2008; Chapman et al. 2009), we chose Homozygosity by locus (HL, (Aparicio et al. 2006) as a metric of diversity within individuals. We used the R package “Genhet” v3.1 (Coulon 2010) to calculate HL. Similarly, because the gut bacterial alpha diversity matric (Shannon) is highly correlated with other matrices (observed ASVs and evenness: R^2^ ≥ 0.8), we used Shannon diversity to estimate diversity within individuals. We examined the relationship between host heterozygosity and bacterial alpha diversity using the *lmer* function in the R package “lme4” v1.1-23 (Bates et al. 2015) to run linear mixed models (LMMs) by setting alpha diversity as the response and heterozygosity as the fixed effect, and collection site as a random effect. The linear mixed model dispersion and residuals were checked with DHARMa v0.3.3.0 (Hartig 2019).

#### Estimation of host genetic, host DNA methylation and gut bacterial diversity and differentiation

We used the R package “phyloseq” v1.32.0 (McMurdie & Holmes 2013) to calculate a Bray Curtis pairwise distance matrices for SNP data, bacterial taxa abundance, and methylation abundance (per locus). Before calculating Bray Curtis distances, We used a Hellinger transformation (Legendre & Gallagher 2001) implemented in the package “microbiome” (Valverde et al. 2014) in R for bacteria and methylation abundance data, which converted absolute abundance to relative abundance. For SNP data, we used TASSEL v5.0 to convert vcf file genotype information into the probability that an allele selected at random at a site is the major allele (e.g. homozygous for major allele = 1.0, homozygous for minor allele = 0.0, and heterozygous genotype = 0.5). We used PCoA analysis through R package “phyloseq” v1.32.0 to visualize data, which is not very sensitive to the influence of double-zeros in the ordination analysis. To compare the diversity of genetic, DNA methylation and gut microbiota between invasion-front and range-core, we calculated the mean and standard deviation (SD) of Bray Curtis distances.

We used the *adonis* command from the package “Vegan” to perform permutational multivariate analysis (the number of permutations = 9999) of variance (perMANOVA) to check whether the cane toad genotype, DNA methylation profile, and bacterial communities from each region were significantly different. We used the command *betadisper* in the package “Vegan” (Oksanen et al. 2019) to check the homogeneity of group variances, an assumption of perMANOVA. After finding significant differences between invasion-front and range-core toads, we performed pairwise comparisons between the six sampling sites using the command *pairwise*.*perm*.*manova* function in “RVAideMemoire” package with the Wilks Lambda (Nath & Pavur 1985) and corrected for multiple testing (Herve 2018) using the Hochberg procedure (Hochberg 1988).

To investigate relationships between host genotype, host DNA methylation and gut bacteria, we used two methods. First, we used a partial Mantel test implemented in the function mantel.partial in the R package “vegan” to compare Bray Curtis distance matrices, while controlling for the effect of geographic distance. After that, we examined the interactions among these three Bray Curtis distance matrices. Then we used LMMs to compare pairwise Bray Curtis distance matrices, accounting for geographic distance (rescaled) and population (invasion-front or range-core) as fixed factors, and individual toad ID as a random factor. We selected the models based on AIC and BIC values and checked their dispersion and residual plots. Each pairwise distance included two individuals: i and j. Each was used in two models: a model with bacterial distance as the response (Bacteria_dist_ij_ ∼ Genotype_dist_ij_ * Methylation_Dist_ij_ + scale(Geographic Distance_ij_) + Population_i_ + Population_j_ + (1|SampleID_i_) + (1|SampleID_j_)); and a model with DNA methylation distance as the response (Methylation_Dist_ij_ ∼ Genotype_dist_ij_ * Bacteria_dist_ij_ + scale(Geographic Distance_ij_) + Population_i_ + Population_j_ + (1|SampleID_i_) + (1|SampleID_j_)). I then used “emmeans” v1.5.4 (V. Lenth et al. 2021) and “ggplot2” v3.3.2 (Wickham 2016) packages in R to visualize the output of the models with an interaction between gut bacterial, host DNA methylation and host genetic distances. To examine the interactions, we visualized how gut bacterial and host DNA methylation varied across different genetic distance classes. We used three representative values to present an infinite set of values with which to fix the continuous genetic distances (UCLA: Statistical Consulting Group). The three representative values of host genetic distances were the mean level of genetic distance, one standard deviation above the mean level of genetic distance, and one standard deviation below the mean level of genetic distance. The slope of the relationship between gut bacteria and host DNA methylation was estimated based on these distance classes, which is a modified version of spotlight analysis (Aiken & West 1991).

## Data Accessibility Statement

All the data and supporting information will be made available online.

1. Supplementary Figures S1.

2. Supplementary Tables S1-2.

The sequence data that support the findings of this study are openly available in NCBI Sequence Read Archive (16S rRNA data accession number: PRJNA670039, msGBS data accession number: PRJNA735013).

## Author Contributions

1. Designed research: JZ, SJZ, LAR, CMRL

2. Performed Research: JZ, KT, LAR, CMRL

3. Analysed data: JZ, KT, SJZ, LAR, CMRL

4. Wrote the paper: JZ, KT, SJZ, LAR, CMRL

## Acknowledgements

This project was supported by the Holsworth Wildlife Research Endowment (Ecological Society of Australia) and Adelaide Graduate Research Scholarship (University of Adelaide) to JZ, and a Scientia Fellowship to LAR (UNSW). CMRL was partially supported by the National Institute of Food and Agriculture, AFRI Competitive Grant Program Accession number 1018617 and National Institute of Food and Agriculture, United States Department of Agriculture, Hatch Program accession number 1020852. KT was supported by the National Institute of Food and Agriculture, United States Department of Agriculture, Hatch Program accession number 1020852. Dr. Eve Slavich from Stats Central in UNSW provided consultations for the statistical analysis. Dr. Alice Russo, and Ms. Rita Kurpiewski contributed to field sample collections. Pastor Jullian Fabres kindly shared the scripts of processing msGBS data. We would like to thank our colleagues from University of Adelaide, University of New South Wales, Macquarie University, University of Kentucky and others who provided insight and expertise that assisted this research.

## Notes

### Competing Interest Statement

The authors have declared no competing interest.

## Bibliography

Aiken LS, West SG. 1991. Multiple regression: Testing and interpreting interactions. Sage Publications, Inc.

Alberdi A, Aizpurua O, Bohmann K, Zepeda-Mendoza ML, Gilbert MTP. 2016. Do vertebrate gut metagenomes confer rapid ecological adaptation? Trends Ecol Evol (Amst). 31:689–699.

Andrews S. 2010. FastQC: A Quality Control Tool for High Throughput Sequence Data [Online]. https://qubeshub.org/resources/fastqc.

Ansari I, Raddatz G, Gutekunst J, Ridnik M, Cohen D, Abu-Remaileh M, Tuganbaev T, Shapiro H, Pikarsky E, Elinav E, et al. 2020. The microbiota programs DNA methylation to control intestinal homeostasis and inflammation. Nat Microbiol. 5:610–619.

Aparicio JM, Ortego J, Cordero PJ. 2006. What should we weigh to estimate heterozygosity, alleles or loci? Mol Ecol. 15:4659–4665.

Ardura A, Zaiko A, Morán P, Planes S, Garcia-Vazquez E. 2017. Epigenetic signatures of invasive status in populations of marine invertebrates. Sci Rep. 7:42193.

Bates D, Maechler M, Bolker B, Walker S. 2015. Fitting Linear Mixed-Effects Models Using lme4. Journal of Statistical Software.

Bell JT, Pai AA, Pickrell JK, Gaffney DJ, Pique-Regi R, Degner JF, Gilad Y, Pritchard JK. 2011. DNA methylation patterns associate with genetic and gene expression variation in HapMap cell lines. Genome Biol. 12:R10.

Blekhman R, Goodrich JK, Huang K, Sun Q, Bukowski R, Bell JT, Spector TD, Keinan A, Ley RE, Gevers D, Clark AG. 2015. Host genetic variation impacts microbiome composition across human body sites. Genome Biol. 16:191.

Bletz MC, Goedbloed DJ, Sanchez E, Reinhardt T, Tebbe CC, Bhuju S, Geffers R, Jarek M, Vences M, Steinfartz S. 2016. Amphibian gut microbiota shifts differentially in community structure but converges on habitat-specific predicted functions. Nat Commun. 7:13699.

Bolnick DI, Snowberg LK, Caporaso JG, Lauber C, Knight R, Stutz WE. 2014. Major Histocompatibility Complex class IIb polymorphism influences gut microbiota composition and diversity. Mol Ecol. 23:4831–4845.

Bolyen E, Rideout JR, Dillon MR, Bokulich NA, Abnet C, Al-Ghalith GA, Alexander H, Alm EJ, Arumugam M, Asnicar F, et al. 2018. QIIME 2: Reproducible, interactive, scalable, and extensible microbiome data science.

Bradbury PJ, Zhang Z, Kroon DE, Casstevens TM, Ramdoss Y, Buckler ES. 2007. TASSEL: software for association mapping of complex traits in diverse samples. Bioinformatics. 23:2633–2635.

Brambilla A, Keller L, Bassano B, Grossen C. 2018. Heterozygosity-fitness correlation at the major histocompatibility complex despite low variation in Alpine ibex (Capra ibex). Evol Appl. 11:631–644.

Callahan BJ, McMurdie PJ, Rosen MJ, Han AW, Johnson AJA, Holmes SP. 2016. DADA2: High-resolution sample inference from Illumina amplicon data. Nat Methods. 13:581–583.

Carja O, MacIsaac JL, Mah SM, Henn BM, Kobor MS, Feldman MW, Fraser HB. 2017. Worldwide patterns of human epigenetic variation. Nat Ecol Evol. 1:1577–1583.

Chapman JR, Nakagawa S, Coltman DW, Slate J, Sheldon BC. 2009. A quantitative review of heterozygosity-fitness correlations in animal populations. Mol Ecol.

Charpentier MJE, Boulet M, Drea CM. 2008. Smelling right: the scent of male lemurs advertises genetic quality and relatedness. Mol Ecol. 17:3225–3233.

Coltman DW, Pilkington JG, Smith JA, Pemberton JM. 1999. Parasite-Mediated Selection against Inbred Soay Sheep in a Free-Living, Island Population. Evolution. 53:1259.

Couch CE, Arnold HK, Crowhurst RS, Jolles AE, Sharpton TJ, Witczak MF, Epps CW, Beechler BR. 2020. Bighorn sheep gut microbiomes associate with genetic and spatial structure across a metapopulation. Sci Rep. 10:6582.

Coulon A. 2010. genhet: an easy-to-use R function to estimate individual heterozygosity. Mol Ecol Resour. 10:167–169.

Cuevas-Sierra A, Ramos-Lopez O, Riezu-Boj JI, Milagro FI, Martinez JA. 2019. Diet, Gut Microbiota, and Obesity: Links with Host Genetics and Epigenetics and Potential Applications. Adv Nutr. 10:S17–S30.

Danecek P, Auton A, Abecasis G, Albers CA, Banks E, DePristo MA, Handsaker RE, Lunter G, Marth GT, Sherry ST, et al. 2011. The variant call format and VCFtools. Bioinformatics. 27:2156–2158.

David I, Canario L, Combes S, Demars J. 2019. Intergenerational transmission of characters through genetics, epigenetics, microbiota, and learning in livestock. Front Genet. 10:1058.

DeSantis TZ, Hugenholtz P, Larsen N, Rojas M, Brodie EL, Keller K, Huber T, Dalevi D, Hu P, Andersen GL. 2006. Greengenes, a chimera-checked 16S rRNA gene database and workbench compatible with ARB. Appl Environ Microbiol. 72:5069–5072.

Dewoody YD, Dewoody JA. 2005. On the estimation of genome-wide heterozygosity using molecular markers. J Hered. 96:85–88.

Dubin MJ, Zhang P, Meng D, Remigereau M-S, Osborne EJ, Paolo Casale F, Drewe P, Kahles A, Jean G, Vilhjálmsson B, et al. 2015. DNA methylation in Arabidopsis has a genetic basis and shows evidence of local adaptation. Elife. 4:e05255.

Eastwood JR, Ribot RFH, Rollins LA, Buchanan KL, Walder K, Bennett ATD, Berg ML. 2017. Host heterozygosity and genotype rarity affect viral dynamics in an avian subspecies complex. Sci Rep. 7:13310.

Edwards RJ, Tuipulotu DE, Amos TG, O’Meally D, Richardson MF, Russell TL, Vallinoto M, Carneiro M, Ferrand N, Wilkins MR, et al. 2018. Draft genome assembly of the invasive cane toad, Rhinella marina. Gigascience. 7.

Estaki M, Pither J, Baumeister P, Little JP, Gill SK, Ghosh S, Ahmadi-Vand Z, Marsden KR, Gibson DL. 2016. Cardiorespiratory fitness as a predictor of intestinal microbial diversity and distinct metagenomic functions. Microbiome. 4:42.

Forstmeier W, Schielzeth H, Mueller JC, Ellegren H, Kempenaers B. 2012. Heterozygosity-fitness correlations in zebra finches: microsatellite markers can be better than their reputation. Mol Ecol. 21:3237–3249.

Goodrich JK, Davenport ER, Beaumont M, Jackson MA, Knight R, Ober C, Spector TD, Bell JT, Clark AG, Ley RE. 2016. Genetic determinants of the gut microbiome in UK twins. Cell Host Microbe. 19:731–743.

Goodrich JK, Waters JL, Poole AC, Sutter JL, Koren O, Blekhman R, Beaumont M, Van Treuren W, Knight R, Bell JT, et al. 2014. Human genetics shape the gut microbiome. Cell. 159:789–799.

Grieneisen L, Muehlbauer AL, Blekhman R. 2020. Microbial control of host gene regulation and the evolution of host-microbiome interactions in primates. Philos Trans R Soc Lond B, Biol Sci. 375:20190598.

Griffiths SM, Harrison XA, Weldon C, Wood MD, Pretorius A, Hopkins K, Fox G, Preziosi RF, Antwis RE. 2018. Genetic variability and ontogeny predict microbiome structure in a disease-challenged montane amphibian. ISME J. 12:2506–2517.

Hartig F. 2019. DHARMa: residual diagnostics for hierarchical (multi-level/mixed) regression models. R package version 02.

Hawes NA, Fidler AE, Tremblay LA, Pochon X, Dunphy BJ, Smith KF. 2018. Understanding the role of DNA methylation in successful biological invasions: a review. Biol Invasions.

Herlemann DP, Labrenz M, Jürgens K, Bertilsson S, Waniek JJ, Andersson AF. 2011. Transitions in bacterial communities along the 2000 km salinity gradient of the Baltic Sea. ISME J. 5:1571–1579.

Herten K, Hestand MS, Vermeesch JR, Van Houdt JKJ. 2015. GBSX: a toolkit for experimental design and demultiplexing genotyping by sequencing experiments. BMC Bioinformatics. 16:73.

Herve M. 2018. Testing and plotting procedures for biostatistics. Package “RVAideMemoire.”

Hochberg Y. 1988. A sharper Bonferroni procedure for multiple tests of significance. Biometrika. 75:800–802.

Kim D, Langmead B, Salzberg SL. 2015. HISAT: a fast spliced aligner with low memory requirements. Nat Methods. 12:357–360.

Kitimu SR, Taylor J, March TJ, Tairo F, Wilkinson MJ, Rodríguez López CM. 2015. Meristem micropropagation of cassava (Manihot esculenta) evokes genome-wide changes in DNA methylation. Front Plant Sci. 6:590.

Krautkramer KA, Kreznar JH, Romano KA, Vivas EI, Barrett-Wilt GA, Rabaglia ME, Keller MP, Attie AD, Rey FE, Denu JM. 2016. Diet-Microbiota Interactions Mediate Global Epigenetic Programming in Multiple Host Tissues. Mol Cell. 64:982–992.

Kreisinger J, Bastien G, Hauffe HC, Marchesi J, Perkins SE. 2015. Interactions between multiple helminths and the gut microbiota in wild rodents. Philos Trans R Soc Lond B, Biol Sci. 370.

Kreznar JH, Keller MP, Traeger LL, Rabaglia ME, Schueler KL, Stapleton DS, Zhao W, Vivas EI, Yandell BS, Broman AT, et al. 2017. Host Genotype and Gut Microbiome Modulate Insulin Secretion and Diet-Induced Metabolic Phenotypes. Cell Rep. 18:1739– 1750.

Kumar H, Lund R, Laiho A, Lundelin K, Ley RE, Isolauri E, Salminen S. 2014. Gut microbiota as an epigenetic regulator: pilot study based on whole-genome methylation analysis. MBio. 5.

Legendre P, Gallagher E. 2001. Ecologically meaningful transformations for ordination of species data. Oecologia. 129:271–280.

Li H, Handsaker B, Wysoker A, Fennell T, Ruan J, Homer N, Marth G, Abecasis G, Durbin R, 1000 Genome Project Data Processing Subgroup. 2009. The Sequence Alignment/Map format and SAMtools. Bioinformatics. 25:2078–2079.

Liebl AL, Schrey AW, Richards CL, Martin LB. 2013. Patterns of DNA methylation throughout a range expansion of an introduced songbird. Integr Comp Biol. 53:351–358.

Lillie M, Shine R, Belov K. 2014. Characterisation of major histocompatibility complex class I in the Australian cane toad, Rhinella marina. PLoS One. 9:e102824.

Luikart G, Pilgrim K, Visty J, Ezenwa VO, Schwartz MK. 2008. Candidate gene microsatellite variation is associated with parasitism in wild bighorn sheep. Biol Lett. 4:228– 231.

Mainguy J, Côté SD, Coltman DW. 2009. Multilocus heterozygosity, parental relatedness and individual fitness components in a wild mountain goat, Oreamnos americanus population. Mol Ecol. 18:2297–2306.

Mayne BT, Leemaqz SY, Buckberry S, Rodriguez Lopez CM, Roberts CT, Bianco-Miotto T, Breen J. 2018. msgbsR: An R package for analysing methylation-sensitive restriction enzyme sequencing data. Sci Rep. 8:2190.

McMurdie PJ, Holmes S. 2013. phyloseq: an R package for reproducible interactive analysis and graphics of microbiome census data. PLoS One. 8:e61217.

McRae AF, Powell JE, Henders AK, Bowdler L, Hemani G, Shah S, Painter JN, Martin NG, Visscher PM, Montgomery GW. 2014. Contribution of genetic variation to transgenerational inheritance of DNA methylation. Genome Biol. 15:R73.

Nath R, Pavur R. 1985. A new statistic in the one-way multivariate analysis of variance. Comput Stat Data Anal. 2:297–315.

Oksanen J, Blanchet FG, Friendly M, Kindt R, Legendre P, McGlinn D, R. Minchin P, O’Hara RB, Simpson GL, Solymos P, et al. 2019. vegan: Community Ecology Package. [place unknown]: vegan: Community Ecology Package.

Penn DJ, Damjanovich K, Potts WK. 2002. MHC heterozygosity confers a selective advantage against multiple-strain infections. Proc Natl Acad Sci USA. 99:11260–11264.

Pereira AC, Bandeira V, Fonseca C, Cunha MV. 2020. Egyptian Mongoose (Herpestes ichneumon) Gut Microbiota: Taxonomical and Functional Differences across Sex and Age Classes. Microorganisms. 8.

Pielou EC. 1966. The measurement of diversity in different types of biological collections. J Theor Biol. 13:131–144.

R Core Team. 2020. R: A language and environment for statistical computing. [place unknown]: R Foundation for Statistical Computing, Vienna, Austria.

Robinson MD, McCarthy DJ, Smyth GK. 2010. edgeR: a Bioconductor package for differential expression analysis of digital gene expression data. Bioinformatics. 26:139–140.

Rollins LA, Richardson MF, Shine R. 2015. A genetic perspective on rapid evolution in cane toads (Rhinella marina). Mol Ecol. 24:2264–2276.

Rothschild D, Weissbrod O, Barkan E, Kurilshikov A, Korem T, Zeevi D, Costea PI, Godneva A, Kalka IN, Bar N, et al. 2018. Environment dominates over host genetics in shaping human gut microbiota. Nature. 555:210–215.

Ryan FJ, Ahern AM, Fitzgerald RS, Laserna-Mendieta EJ, Power EM, Clooney AG, O’Donoghue KW, McMurdie PJ, Iwai S, Crits-Christoph A, et al. 2020. Colonic microbiota is associated with inflammation and host epigenomic alterations in inflammatory bowel disease. Nat Commun. 11:1512.

Sarma RR, Edwards RJ, Crino OL, Eyck HJF, Waters PD, Crossland MR, Shine R, Rollins LA. 2020. Do epigenetic changes drive corticosterone responses to alarm cues in larvae of an invasive amphibian? Integr Comp Biol. 60:1481–1494.

Schubert M, Lindgreen S, Orlando L. 2016. AdapterRemoval v2: rapid adapter trimming, identification, and read merging. BMC Res Notes. 9:88.

Selechnik D, Richardson MF, Shine R, Brown GP, Rollins LA. 2019. Immune and environment-driven gene expression during invasion: An eco-immunological application of RNA-Seq. Ecol Evol. 9:6708–6721.

Selechnik D, Richardson MF, Shine R, DeVore JL, Ducatez S, Rollins LA. 2019. Increased adaptive variation despite reduced overall genetic diversity in a rapidly adapting invader. Front Genet. 10:1221.

Shannon CE. 1948. A mathematical theory of communication. Bell System Technical Journal. 27:379–423.

Sheldon EL, Schrey A, Andrew SC, Ragsdale A, Griffith SC. 2018. Epigenetic and genetic variation among three separate introductions of the house sparrow (Passer domesticus) into Australia. Royal Society Open Science.

Smith CCR, Snowberg LK, Gregory Caporaso J, Knight R, Bolnick DI. 2015. Dietary input of microbes and host genetic variation shape among-population differences in stickleback gut microbiota. ISME J. 9:2515–2526.

Stilling RM, Dinan TG, Cryan JF. 2014. Microbial genes, brain & behaviour - epigenetic regulation of the gut-brain axis. Genes Brain Behav. 13:69–86.

Suzuki TA. 2017. Links between Natural Variation in the Microbiome and Host Fitness in Wild Mammals. Integr Comp Biol. 57:756–769.

Suzuki TA, Phifer-Rixey M, Mack KL, Sheehan MJ, Lin D, Bi K, Nachman MW. 2019. Host genetic determinants of the gut microbiota of wild mice. Mol Ecol. 28:3197–3207.

Tong Q, Cui L-Y, Du X-P, Hu Z-F, Bie J, Xiao J-H, Wang H-B, Zhang J-T. 2020. Comparison of Gut Microbiota Diversity and Predicted Functions Between Healthy and Diseased Captive Rana dybowskii. Front Microbiol. 11:2096.

UCLA: Statistical Consulting Group. Decomposing, probing, and plotting interactions in R [Internet]. [cited 2021 Apr 19]. Available from: https://stats.idre.ucla.edu/r/seminars/interactions-r/#s3

Uren Webster TM, Consuegra S, Hitchings M, Garcia de Leaniz C. 2018. Interpopulation variation in the atlantic salmon microbiome reflects environmental and genetic diversity. Appl Environ Microbiol. 84.

Ussar S, Griffin NW, Bezy O, Fujisaka S, Vienberg S, Softic S, Deng L, Bry L, Gordon JI, Kahn CR. 2015. Interactions between Gut Microbiota, Host Genetics and Diet Modulate the Predisposition to Obesity and Metabolic Syndrome. Cell Metab. 22:516–530.

V. Lenth R Buerkner P, Herve M, Love J, Riebl H, Singmann H. 2021. emmeans: Estimated Marginal Means, aka Least-Squares Means. [R package].

Valverde A, Makhalanyane TP, Cowan DA. 2014. Contrasting assembly processes in a bacterial metacommunity along a desiccation gradient. Front Microbiol. 5:668.

Velando A, Barros Á, Moran P. 2015. Heterozygosity-fitness correlations in a declining seabird population. Mol Ecol. 24:1007–1018.

Wen C, Yan W, Sun C, Ji C, Zhou Q, Zhang D, Zheng J, Yang N. 2019. The gut microbiota is largely independent of host genetics in regulating fat deposition in chickens. ISME J. 13:1422–1436.

Wickham H. 2016. ggplot2: Elegant Graphics for Data Analysis. Springer-Verlag New York. [R package].

Wu J, Zhao Y, Wang X, Kong L, Johnston LJ, Lu L, Ma X. 2020. Dietary nutrients shape gut microbes and intestinal mucosa via epigenetic modifications. Crit Rev Food Sci Nutr.:1–15.

Xu F, Fu Y, Sun T-Y, Jiang Z, Miao Z, Shuai M, Gou W, Ling C-W, Yang J, Wang J, et al. 2020. The interplay between host genetics and the gut microbiome reveals common and distinct microbiome features for complex human diseases. Microbiome. 8:145.

Yu D-H, Gadkari M, Zhou Q, Yu S, Gao N, Guan Y, Schady D, Roshan TN, Chen M-H, Laritsky E, et al. 2015. Postnatal epigenetic regulation of intestinal stem cells requires DNA methylation and is guided by the microbiome. Genome Biol. 16:211.

Zhang C, Zhang M, Wang S, Han R, Cao Y, Hua W, Mao Y, Zhang X, Pang X, Wei C, et al. 2010. Interactions between gut microbiota, host genetics and diet relevant to development of metabolic syndromes in mice. ISME J. 4:232–241.

Zhou J, Nelson TM, Rodriguez Lopez C, Zhou SJ, Ward-Fear G, Stuart KC, Rollins LA. 2020. Microbial function is related to behavior of an invasive anuran. BioRxiv.

