## supplementary figure for "Genetic similarity enhances the strength of the relationship between gut bacteria and host DNA methylation"

***Supplementary Material***

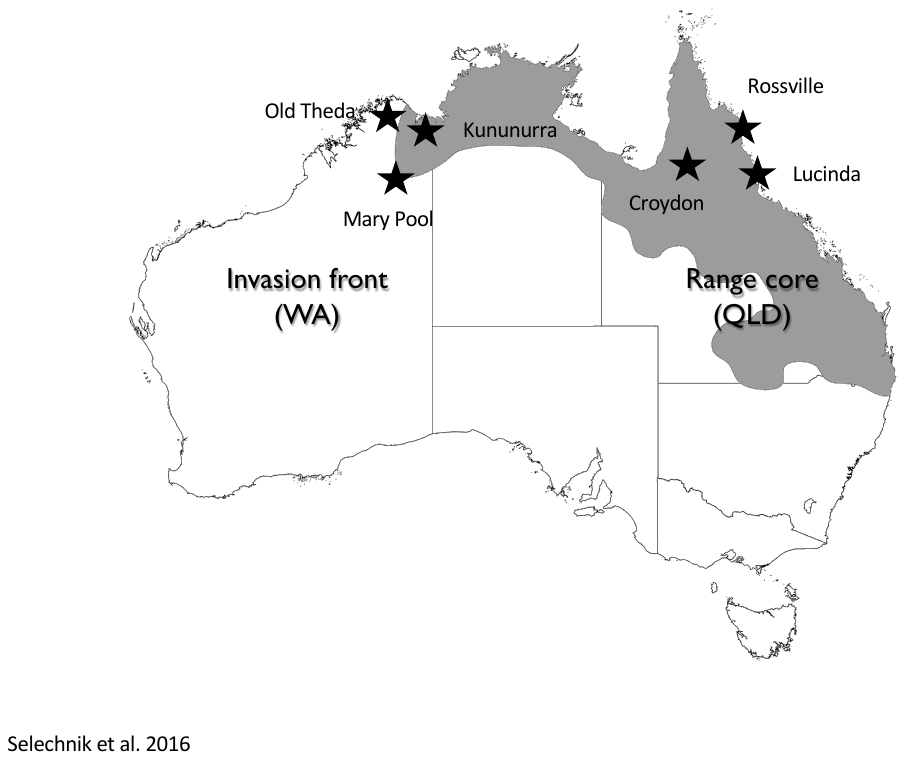

**Figure S1. Sampling sites.** Grey area indicates the geographic distribution of cane toads in Australia at the time of sampling. Female individuals (n=10 per site) from three sites near the invasion-front (Kununurra 15.776566° S, 128.744293° E, Old Theda 14.790795° S, 126.497624° E, and Mary Pool 18.72528° S, 126.870096° E), and three sites near the range-core (Rossville 15.697069° S, 145.254385° E, Croydon 18.207536° S, 142.245702° E, and Lucinda 18.530149° S, 146.331264° E) were collected in November 2018 and December 2018 respectively. Adapted from Selechnik et al., (2017).
